# The roseoloviruses downregulate the protein tyrosine phosphatase PTPRC (CD45)

**DOI:** 10.1101/2020.09.29.318709

**Authors:** Melissa L. Whyte, Kelsey Smith, Amanda Buchberger, Linda Berg Luecke, Lidya Handayani Tjan, Yasuko Mori, Rebekah L Gundry, Amy W. Hudson

**Affiliations:** Department of Microbiology and Immunology, Medical College of Wisconsin, Milwaukee, WI; Department of Biochemistry, Medical College of Wisconsin, Milwaukee, WI; Division of Clinical Virology, Kobe University Graduate School of Medicine, Kobe, Japan; CardiOmics Program, Center for Heart and Vascular Research; Division of Cardiovascular Medicine; and Department of Cellular and Integrative Physiology, University of Nebraska Medical Center, Omaha, NE

## Abstract

Like all herpesviruses, the roseoloviruses (HHV6A, -6B, and -7) establish lifelong infection within their host, requiring these viruses to evade host anti-viral responses. One common host-evasion strategy is the downregulation of host-encoded, surface-expressed glycoproteins. Roseoloviruses have been shown to evade host the host immune response by downregulating NK-activating ligands, MHC class I, and the TCR/CD3 complex. To more globally identify glycoproteins that are differentially expressed on the surface of HHV6A-infected cells, we performed cell surface capture of N-linked glycoproteins present on the surface of T cells infected with HHV6A, and compared these to proteins present on the surface of uninfected T cells. We found that the protein tyrosine phosphatase CD45 is downregulated in T cells infected with HHV6A. We also demonstrated that CD45 is similarly downregulated in cells infected with HHV-7. CD45 is essential for signaling through the T cell receptor and as such, is necessary for developing a fully functional immune response. Interestingly, the closely related β-herpesviruses human cytomegalovirus (HCMV) and murine cytomegalovirus (MCMV) have also separately evolved unique mechanisms to target CD45. While HCMV and MCMV target CD45 signaling and trafficking, HHV6A acts to downregulate CD45 transcripts.

**Importance:** Human herpesviruses-6 and -7 infect essentially 100% of the world’s population before the age of 5 and then remain latent or persistent in their host throughout life. As such, these viruses are among the most pervasive and stealthy of all viruses. Host immune cells rely on the presence of surface-expressed proteins to identify and target virus-infected cells. Here, we investigated the changes that occur to proteins expressed on the cell surface of T cells after infection with human herpesvirus-6A. We discovered that HHV-6A infection results in a reduction of CD45 on the surface of infected cells. Targeting of CD45 may prevent activation of these virus-infected T cells, possibly lengthening the life of the infected T cell so that it can harbor latent virus.

## Introduction

Human Herpesvirus-6A (HHV6A) is a human-specific, T cell-tropic β-herpesvirus that is most closely related to the other members of the roseolovirus genus, HHV-6B and HHV-7, as well as human cytomegalovirus (HCMV). Primary infections with HHV-6 and -7 usually occur before the age of three and are often characterized by a high fever (1, 2). HHV-6 and -7 infect over 90% of the population, and like other herpesviruses, HHV-6 and -7 remain latent or establish lifelong infections in their hosts. As such, these viruses are among the most pervasive and stealthy of all viruses; they must necessarily excel at escaping immune detection throughout the life of the host, yet little is known about how these viruses so successfully escape host defenses.

The ability of host immune cells to detect virus-infected cells is largely dependent on interactions between proteins expressed on the surface of immune cells and those expressed on the surface of their targets. Viruses have necessarily evolved to alter the expression of these cell-surface-expressed proteins to evade detection by the host immune system. The herpesviruses, particularly the cytomegaloviruses, have long been known to employ devious strategies to interfere with expression of host-encoded surface-expressed proteins. For example, most herpesviruses, including HHV-6 and -7, interfere with antigen presentation by downregulating class I major histocompatibility complex (MHC) molecules from the surface of infected cells (4-13). Additionally, HCMV, the most closely-related virus to the roseoloviruses, encodes at least 5 different gene products and a miRNA that all participate in downregulation of natural killer (NK)-activating ligands, all in an effort to prevent natural killer cells from identifying and killing infected cells (14-18). We have found that HHV6A, -6B, and -7 each encode a single gene product, U21, that acts as a multifunctional transmembrane glycoprotein to downregulate not only multiple NK activating ligands, but also most class I MHC alleles (12, 19-21).

Herpesviruses also target surface-expressed proteins to hinder the ability of immune cells to perform their effector function. For example, herpesviruses inhibit T cell function through the downregulation of co-stimulatory ligands like B7 and CD40, and upregulation of co-inhibitory ligands such as PD-L1 and galectin-9 (22-32). Herpesviruses also downregulate adhesion molecules, such as I-CAM, V-CAM, PECAM and ALCAM, which can physically disrupt formation of the immune synapse and inhibit T cell activation (33-37).

Since most herpesviruses are not T cell-tropic, changes in surface expression occur in infected cells, and infected cells then exert their immunosuppressive effects through interaction with uninfected T cells. But what happens to the surface of a T cell when it is infected with a herpesvirus? The T cell-tropic roseoloviruses allow us to explore and learn from alterations to the glycoprotein landscape on the surface of herpesvirus-infected T cells. Herein, we applied a mass spectrometry strategy to identify and quantify N-linked glycoproteins on the cell surface of HHV6A-infected T cells to yield the first insights into how virus infection induces alterations in the cell surface landscape of host cells. We discovered that the protein tyrosine phosphatase CD45 (PTPRC) is downregulated from the surface of HHV6A-infected T cells. CD45 is expressed on the surface of all nucleated cells of hematopoietic origin, where its activity is critical for the proper function of immune cells (reviewed in (39-42)). The phosphatase activity of CD45 is required for successful signaling through the TCR (43-45), and as such, the downregulation of CD45 could be an attractive strategy for viruses to impair T cell signaling.

## Results

### Cell-surface capture of CD45 in HHV6A-infected T cells

To identify host-encoded surface-expressed proteins targeted by HHV6A, we performed cell surface capture (CSC), a chemoproteomic approach for the specific identification of cell surface *N*-glycoproteins (46-50). Using CSC and bioinformatic analysis tools, we compared N-linked glycoproteins identified on the surface of T cells infected with HHV6A to those identified on the surface of uninfected T cells (3, 38). To minimize biological variability, we performed CSC in HHV6A-infected cell line. HHV6A infects only a small number of cultured cell lines, and for these studies, we used JJhan cells, a CD4+ T cell line variant of Jurkat cells (51).

We identified 605 cell surface N-linked glycoproteins on the surface of infected and uninfected JJhan cells, including 594 human and 11 viral proteins. Strikingly, we identified 47 unique peptides derived from the protein tyrosine phosphatase PTPRC (CD45), spanning all 22 N-linked glycosylation sites (Fig. 1a). CD45-derived peptides were consistently less abundant on the surface of HHV6A-infected JJhan cells across multiple biological and technical replicates (Fig. 1b-d), suggesting that CD45 is downregulated from the surface of HHV6A-infected JJhan cells.

**Figure 1.**
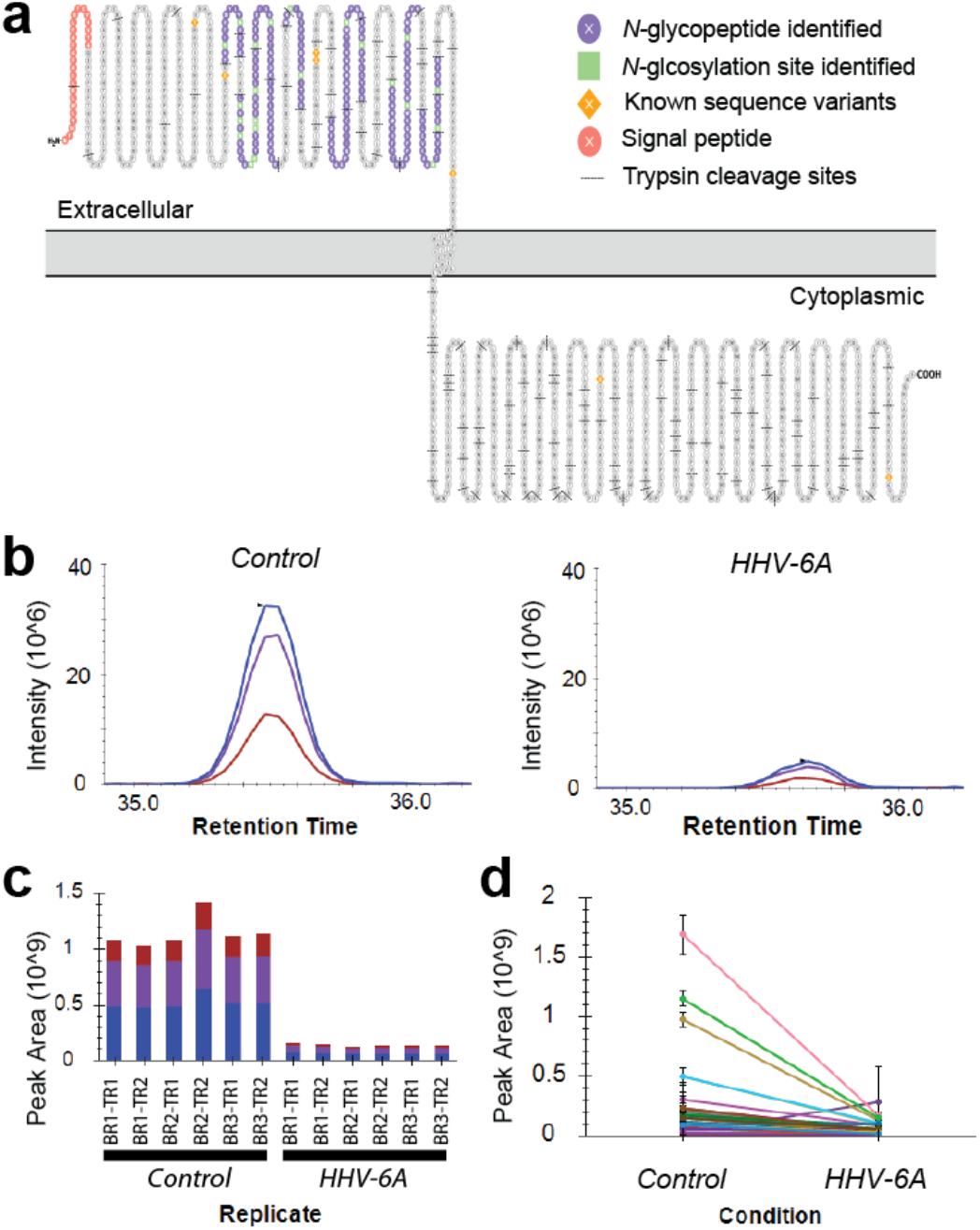
Mass spectrometry data for CD45. (a) Graphical representation of CD45 protein with extracellular N-linked glycopeptides identified by CSC indicated. This approach specifically enriches and identifies N-glycopeptides from the extracellular domain of cell surface proteins. Image generated using Protter (3). (bd) Label-free quantitation data for CD45. Representative peaks of a peptide (YAnITVDYLYNK, where n is the site of N-glycosylation) detected across individual replicate analyses (b) and three replicate analyses (including technical replicates) (c) for each sample group, showing CD45 is more abundant in control compared to infected cells. Each color (blue, purple, and red) are associated with the peak area from the [M]+0, [M]+1, and [M]+2 isotopic peaks, respectively. [M] is the monoisotopic mass. (d) Summary of peak areas for all 67 peptide observations for CD45 across all technical and biological replicates. Each line is a different peptide. BR: biological replicate; TR: technical replicate. Results shown in b-d were generated using Skyline (38).

### Localization and steady-state expression of CD45 in HHV6A-infected T cells

To independently validate the downregulation of CD45 observed in our mass spectrometry experiments, we examined surface-expression of CD45 in HHV6A-infected cells by flow cytometry. We infected JJhan cells with a recombinant, bacterial artificial chromosome (BAC)-derived HHV6A virus encoding soluble green fluorescent protein (GFP), which allowed us to identify actively-infected cells (52). In uninfected JJhan cells, we observed a single population of CD45-expressing cells, while in HHV6A-infected JJhan cells, we observed reduced surface expression of CD45 (Fig. 2a), consistent with our mass spectrometry finding that the presence of CD45 is diminished on the surface of HHV6A-infected cells.

**Figure 2.**
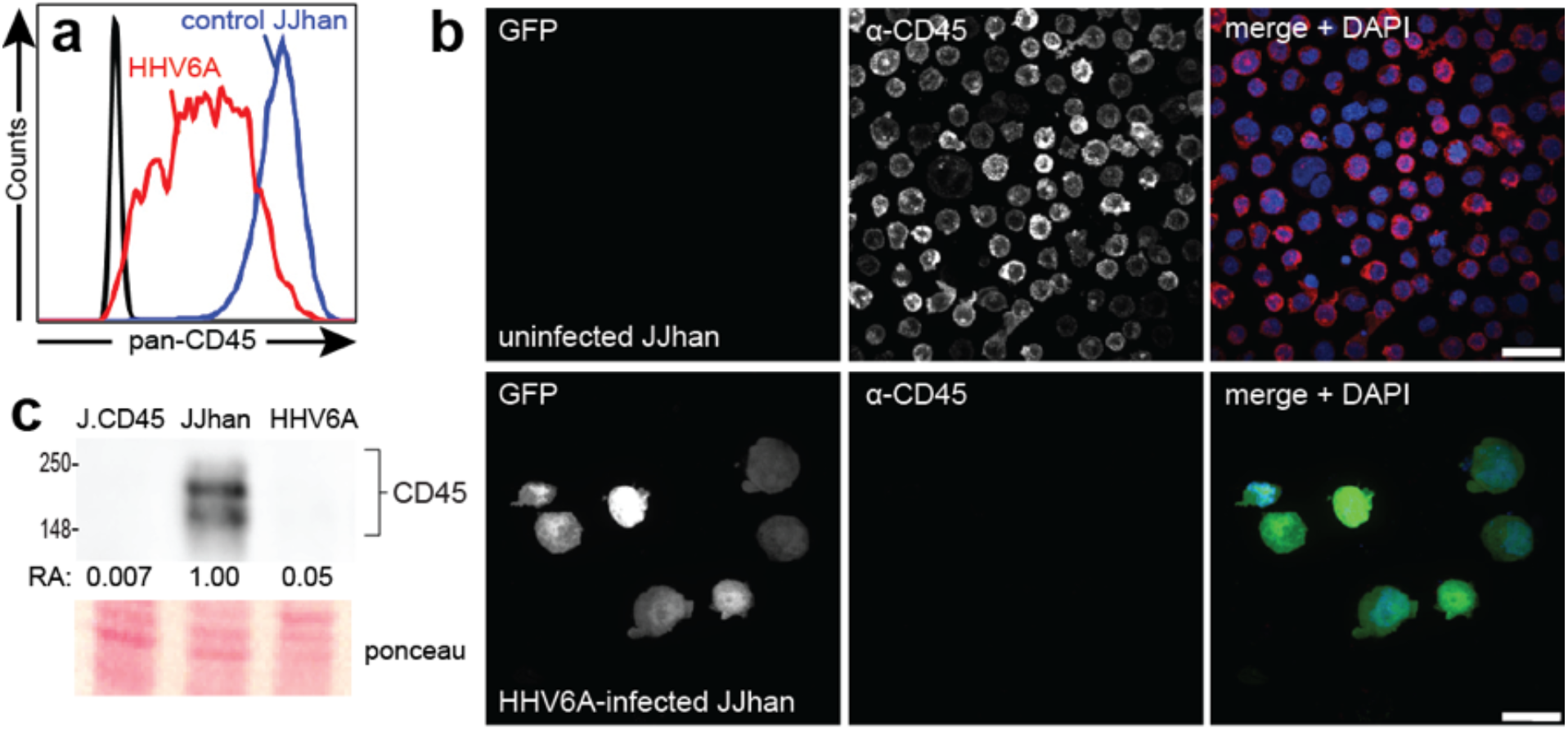
CD45 is downregulated in HHV6A-infected cells. (a) Flow cytometric analysis of HHV6A-infected JJhan cells (red) and uninfected JJhan cells (blue). HHV6A-infected cells were gated on as GFP+. Live cells were labeled with an antibody directed against CD45. (b) Confocal immunofluorescence microscopy images showing uninfected JJhan cells (top panels) and HHV6A-infected (GFP+) JJhan cells (bottom panels) that were fixed, permeabilized, and labeled with an antibody directed against CD45. Images were taken at 100X magnification and are shown as maximum intensity projections. Scale bar = 20 μm. (c) Immunoblot analysis of CD45 in whole cell lysates generated from J.CD45 cells (negative control), uninfected JJhan cells, and HHV6A-infected JJhan cells. Protein was normalized to total protein for each lane. RA: relative abundance. Abundance is calculated relative to uninfected JJhan cells. Ponceau shows total protein loaded per lane.

CD45 is expressed as multiple different isoforms in human cells (53-55). To determine whether all isoforms of CD45 are downregulated in HHV6A-infected JJhan cells, we labeled cells with antibodies directed against the CD45 isoforms commonly expressed in Jurkat cells, the parental cell line of JJhan cells (56). CD45-RO, -RB, and -RC were all downregulated in HHV6A-infected JJhan cells relative to uninfected JJhan cells (Fig. S1), suggesting the downregulation of CD45 from the surface HHV6A-infected JJhan cells is not specific to any one CD45 isoform.

Viruses often reroute the intracellular trafficking of surface-expressed proteins from the cell surface to alter host cell biology. For example, the HHV6A U24 gene product downregulates the T-cell receptor (TCR)/CD3 by stimulating endocytosis of the receptor resulting in relocalization of TCR/CD3 from the cell surface to endosomes, and U21 reroutes class I MHC molecules to lysosomes (13, 57). We therefore sought to determine whether the reduction in surface-expressed CD45 observed in HHV6A-infected cells was the result of a redistribution of CD45 within the cell. To examine the localization of CD45, we performed immunofluorescence microscopy of permeabilized cells labeled with an antibody directed against human CD45. As expected, CD45 was localized to the plasma membrane in uninfected JJhan T cells. In HHV6A-infected JJhan cells, however, we observed a striking disappearance of CD45 labeling (Fig. 2b).

To better quantify the downreguation of CD45 in HHV6A-infected JJhan cells, we next examined steady-state levels of CD45 protein by immunoblot analysis of whole-cell lysates generated from uninfected or HHV6A-infected JJhan cells. Consistent with our immunofluorescence microscopy data (Fig. 2b), immunoblot analysis showed a 95% reduction in CD45 protein in HHV6A-infected JJhan cells relative to uninfected JJhan cells (Fig. 2c). Taken together, these results demonstrate that CD45 is not only downregulated from the surface of HHV6A-infected JJhan cells but is also depleted from HHV6A-infected cells.

### CD45 expression in HHV7-infected T cells

The HHV6A genome is almost entirely co-linear with the two other roseolovirus genomes HHV-6B and HHV-7 (58, 59). As such, we reasoned that HHV-6B and HHV-7 infection may also result in CD45 downregulation. To determine whether CD45 is downregulated in HHV7-infected cells, we evaluated the localization of CD45 in HHV7-infected SupT1 cells by immunofluorescence microscopy. Since a recombinant GFP-containing BAC containing the coding sequence of HHV-7 is not yet available, we identified HHV7-infected cells using an antibody directed against the U21 gene product from HHV-7 (Fig. 3, green). Similar to cells infected with HHV6A (Fig. 2b), HHV7-infected SupT1 cells showed reduced CD45 labeling (Fig. 3, bottom panels as compared to uninfected SupT1 cells (Fig. 3, top panels), suggesting that the ability to downregulate CD45 is common to all roseoloviruses.

**Figure 3.**
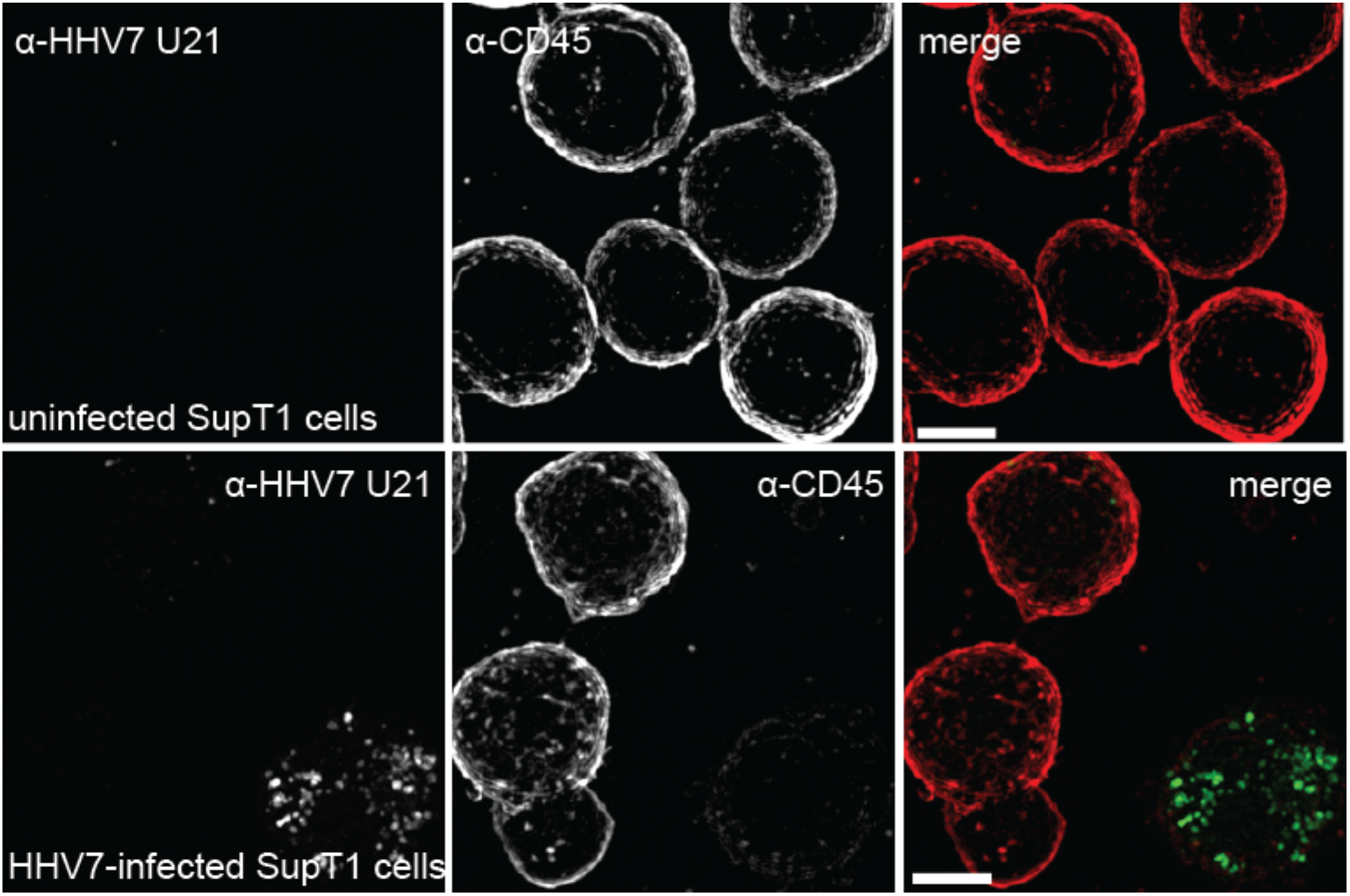
CD45 is downregulated in HHV7-infected T cells. Structured Illumination 3-dimensional microscopy images showing uninfected control T cells (top panels) and HHV7-infected T cell (bottom panels). Fixed and permeabilized cells were labeled with an antibody directed against the U21 open reading frame (ORF) from HHV7 (green) and an antibody directed against human CD45 (red). Images were taken at 100X and are shown as maximum intensity projections. Scale bar = 5 μm.

### CD45 protein stability in HHV6A-infected T cells

Viruses often employ the strategy of degrading host proteins involved in the host response to virus infection. For example, the murine cytomegalovirus m42 gene product induces internalization and subsequent proteasomal degradation of CD45 (60). To further investigate the mechanism by which CD45 is downregulated in HHV6A-infected JJhan cells, we examined the stability of CD45 in the presence of proteasomal and lysosomal protease inhibitors.

Lysosomal protease function is compromised at a more neutral pH, thus stabilization of a protein in the presence of the weak base ammonium chloride (NH_4_Cl) would suggest the involvement of lysosomal proteases in its turnover. Likewise, stabilization of a protein in the presence of the peptide-aldehyde MG-132, which selectively inhibits proteolytic activity of the 26S proteasome, would suggest involvement of proteasomal proteases in the turnover of that protein. We separately inhibited lysosomal and proteasomal protein degradation, treating cells with NH_4_Cl or MG-132, and assessed steady-state CD45 protein levels by immunoblot analysis. In cells treated with a DMSO vehicle control, we observed a 95% reduction in CD45 abundance in HHV6A-infected cells (Fig 4a), which is similar to what we observed in untreated cells (Fig. 2c), suggesting the dimethyl sulfoxide (DMSO) vehicle does not affect degradation of CD45. We observed a similar reduction in CD45 in cells treated with MG-132 or NH4Cl (Fig. 4a).

**Figure 4.**
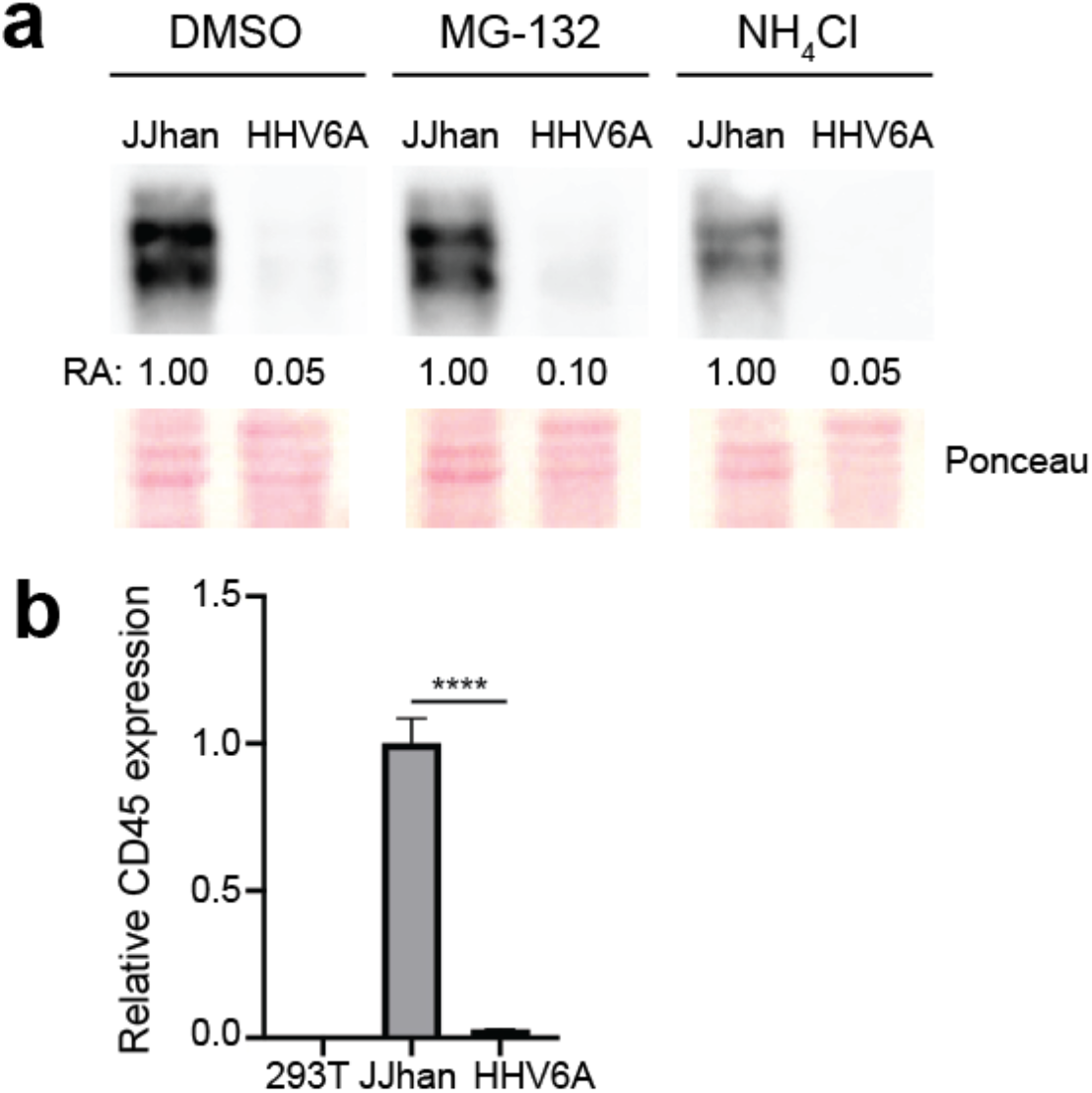
CD45 is downregulated at the RNA level in HHV6A-infected T cells. (a) Immunoblot analysis showing steady-state abundance of CD45 protein in uninfected JJhan cells, or HHV6A-infected cells under different conditions (DMSO vehicle control, MG-132, or NH4Cl treatment for 19 hours). Equal protein was loaded in each lane, and CD45 abundance was calculated relative to uninfected JJhan cells for each treatment. RA = relative abundance. Ponceau-stained lanes visually illustrate equal protein loading. (b) CD45 RNA abundance in 293T cells (negative control), uninfected JJhan cells, or HHV6A-infected cells. CD45 RNA abundance is calculated relative to a cellular control gene and shown here relative to uninfected JJhan cells. Data shown is from a single representative biological replicate, performed in technical triplicate for each sample, mean +/- SD. **** = p < 0.0001.

We also examined localization of CD45 in HHV6A-infected cells treated with MG-132 or NH_4_Cl by immunofluorescence microscopy. As expected, we observed surface-localization of CD45 in untreated, uninfected JJhan cells (Fig. S2, panel a). We also observed surface-localization of CD45 in uninfected JJhan cells treated with a DMSO vehicle, MG-132, or NH4Cl (Fig. S2, panels b, c, and d, respectively), suggesting CD45 localization is not affected by treatment with MG-132 or NH_4_Cl outside the context of an HHV6A infection. Consistent with the immunoblot data, there was little to no CD45 labeling present in untreated HHV6A-infected JJhan cells (Fig. S2, panel e). Similarly, there was little to no CD45 labeling in HHV6A-infected treated with a DMSO vehicle, MG-132, or NH_4_Cl (Fig. S2, panels f, g, & h, respectively). Taken together, these results suggest that the downregulation of CD45 in HHV6A-infected JJhan cells occurs by some method other than protein degradation.

### CD45 gene expression in HHV6A-infected T cells

Because CD45 was not stabilized in the presence of proteasomal or lysosomal inhibitors, we hypothesized that CD45 was downregulated at the transcript level in HHV6A-infected T cells. To test this hypothesis, we quantified CD45 mRNA in uninfected and HHV6A-infected JJhan cells by quantitative reverse transcriptase-polymerase chain reaction (qRT-PCR). CD45 mRNA levels were greatly reduced in JJhan cells infected with HHV6A as compared to uninfected JJhan cells (Fig. 4b), suggesting that CD45 transcripts are downregulated in HHV6A-infected JJhan cells.

## Discussion

Herpesviruses, as life-long pathogens, are especially masterful at reprogramming host cells to create a more hospitable environment. Perhaps the greatest challenge to a virus is the detection and killing of its host cell by immune cells. As discussed, the herpesviruses encode multiple gene products that alter host cell biology to prevent the identification of infected cells. These measures are not entirely sufficient, however, and viruses also strategize to target immune cells to inhibit their functional capacity. One way viruses can inhibit these processes is by targeting the proteins that are important for immune cell function, such as the protein tyrosine phosphatase CD45.

CD45 is expressed on the surface of all nucleated cells of hematopoietic origin, where its activity is critical for the proper function of immune cells (reviewed in (39-42)). In T cells, CD45’s primary substrates are Src family kinases (61-63). CD45 dephosphorylates the inhibitory phosphotyrosine residue on the Src kinase Lck, leaving the kinase in a ‘primed’ state, so that it can be activated through T cell receptor (TCR) signaling (62). The phosphatase activity of CD45 is required for successful signaling through the TCR (43-45), and as such, the downregulation of CD45 could be an attractive strategy for viruses to inhibit T cell signaling.

Here we describe the downregulation of CD45 by two roseoloviruses, HHV6A and HHV-7. Expression of CD45 is markedly reduced in HHV6A-infected JJhan cells and HHV7-infected SupT1 cells. While we do not yet fully understand the functional consequences of CD45 downregulation in roseolovirus-infected cells, we can gather clues from three other viruses also known to target CD45: human cytomegalovirus (HCMV), human adenovirus, and murine cytomegalovirus (MCMV).

CD45 is targeted by HCMV pUL11 (64). The extracellular domain of pUL11, which is expressed on the surface of HCMV-infected epithelial cells or fibroblasts, interacts with CD45 on nearby uninfected T cells. The interaction between pUL11 and CD45 inhibits the phosphatase activity of CD45, which in turn impairs TCR signaling and ultimately, T cell proliferation (64, 65). Inhibition of CD45 by pUL11 also results in an increase in production of the anti-inflammatory cytokine IL-10 (65). In HHV6A infection, the concentration of secreted IL-10 protein was shown to be increased during HHV6A infection at timepoints up to 72 hours post-infection (66, 67). Since the inhibition of CD45 during HCMV infection results in an increase in IL-10 production (65), it is tempting to speculate that the downregulation of CD45 we observe in HHV6A-infected T cells may be involved in the increase in IL-10 levels shown to occur during HHV6A infection.

Another virus that devotes an open reading frame to the downregulation of CD45 is Adenovirus19a (Ad19a). Ad19a E3/49K protein is cleaved to a secreted form (sec49K) that interacts with CD45 on nearby uninfected NK cells and T cells. Sec49K-mediated inhibition of CD45 suppresses T cell activation and signaling, resulting in diminished production of the anti-viral cytokine IFN-γ, and inhibition of NK cell activation (68). After HHV6A infection of stimulated peripheral blood mononuclear cells, the production of IFN-γ was also reported to be reduced (66). Since inhibition of CD45 function during adenovirus infection results in a decrease in IFN-γ production (68), it is possible that the decrease reported in IFN-γ production during HHV6A infection may occur as a result of CD45 downregulation in infected T cells.

Unlike HCMV pUL11 and Ad19a sec49K, which act extracellularly to inhibit CD45, MCMV encodes a protein, m42, that induces the internalization and degradation of CD45 within MCMV-infected macrophages (60). MCMV m42 acts as an adaptor or activator of HECT3 E3 ubiquitin ligases, and through ubiquitination, m42 marks CD45 for lysosomal degradation (60). The functional outcome of m42-mediated downregulation of CD45 in MCMV-infected macrophages is unclear. HHV6A-mediated downregulation of CD45 is similar to MCMV-mediated downregulation of CD45, in that these viruses downregulate CD45 from within infected cells, as opposed to acting on CD45 *in trans* on the surface of nearby uninfected cells. As yet, the functional outcome of downregulating CD45 within either of these infected immune cells remains elusive.

The roseolovirus genomes lack positional or functional homologs of MCMV m42 or HCMV pUL11, and unlike HCMV, MCMV, and Ad19a, HHV6A infection results in the dramatic transcriptional downregulation of CD45. HHV6A therefore downregulates CD45 by a novel mechanism. The separate evolution of four unique mechanisms to target a single host protein strongly suggests that CD45 is an important viral target, though its impact is unclear: how might the roseoloviruses benefit from the downregulation of CD45 in infected T cells? As discussed above, while the influence of CD45 downregulation on cytokine production would certainly benefit HHV6A, it is important to note that HHV6A preferentially infects T cells. Since it takes days to mount a virus-specific T cell response, during a primary infection, the T cells that HHV6A infects are not likely to be HHV6A-specific. Therefore, downregulation of CD45 in infected T cells would not directly impair activation of T cells responding to HHV6A infection. Instead, downregulation of CD45 may be a means to inhibit activation of the HHV6A-infected T cell, possibly preventing activation-induced cell death, and creating a host cell environment conducive to harboring latent virus. Future work is focused on identification of the HHV6A gene products involved in the transcriptional regulation of CD45 and exploring the functional consequences of CD45 downregulation in roseolovirus-infected T cells.

## Materials & Methods

### Cell lines and viruses

JJhan, ND10^depl^ JJhan, and J.CD45 T cells were cultured in RPMI-1640 medium (ThermoFisher Scientific, Waltham, MA) supplemented with 5% fetal bovine serum (FBS) and 2 mM L-glutamine. JJhan cells depleted of ND10 (ND10^depl^ JJhan cells) were the kind gift of Dr. Benedikt Kaufer (Freie Universität, Berlin, Germany) (69). J.CD45 cells (CD45 -negative Jurkat cells) were kindly provided by Dr. Arthur Weiss (UCSF, San Francisco, CA) (70). 293T cells were cultured in Dulbecco’s modified Eagle medium (DMEM) (Invitrogen, Carlsbad, CA) supplemented with 5% FBS, 5% newborn calf serum, and 2 mM L-glutamine. The HHV6A virus used in these studies is a recombinant HHV6A virus (strain U1102) generated from a bacterial artificial chromosome (BAC) containing the HHV6A genome with GFP inserted between the U53 and U54 ORFs (71). Infectious HHV6A virus was generated by electroporating HHV6A-GFP BAC DNA into JJhan T cells. HHV6A virus was propagated by mixing uninfected ND10-depleted Jhan cells with infected JJhan cells once HHV6A-infected JJhan cells were >80% GFP+. HHV6A-infected and uninfected JJhan cells as well as J.CD45 cells were stimulated with 3.75 ug/ml Phytohemagglutinin (PHA-P) and 9 ug/ml hydrocortisone. HHV6A-infected cells used for experiments in this study were ≥70% GFP+ at the time of harvest. SupT1 cells were cultured in RPMI-1640 medium supplemented with 2.5% FBS and 2 mM L-Glutamine. HHV-7 infection (strain SB) was performed by co-culture of HHV7-infected SupT1 cells with uninfected SupT1 cells.

### Antibodies & reagents

The monoclonal anti-human CD45 antibody (clone S5-Z6, Santa Cruz Biotechnology, Dallas, TX) was used for flow cytometry (1 ug/ 1 x10^6^ cells), immunofluorescence microscopy (1:50), and immunoblotting (1:50). The monoclonal anti-CD45 antibody (clone MEM-28, Millipore-Sigma, St. Louis, MO) was used for immunofluorescence microscopy (1:200). The polyclonal antibody MCW62 (U21-N) was raised against the N-terminus of HHV-7 U21 (1:400) (72). AlexaFluor-405, -488, -594, and -647-conjugated goat-anti mouse and rabbit secondary antibodies were used at dilutions recommended by the manufacturer (ThermoFisher Scientific). Chemicals were purchased from Sigma Millipore unless otherwise noted.

### Identification of cell surface N-glycoproteins

Cell Surface Capture (CSC) was applied to control JJhan and HHV6A-infected JJhan T cells (10 million cells per replicate, three biological replicates per condition). Briefly, extracellular glycans on intact cells were oxidized using sodium meta-periodate, and the resulting aldehydes were labelled with biocytin hydrazide to form a ‘handle’ for enrichment. Cells were then lysed, proteins enzymatically digested, and biotinylated glycopeptides were enriched using streptavidin beads. *N*-glycopeptides were then selectively released by Peptide-N-Glycosidase F (PNGase:F) and analysed by mass spectrometry. Here, CSC was performed as previously described in detail (47-50), with the exception that glycopeptide enrichment and bead washing was performed using an epMotion 5073m (Eppendorf, Hamburg, Germany). For each sample, 750 μg of total peptide was diluted in binding buffer (80 mM sodium phosphate, 2 M NaCl, 0.2% Tween 20, pH 7.8) and incubated with 100μL of GenScript Streptavidin MagBeads (GenScript, Piscataway, NJ) for 1 h with mixing. Beads were then washed sequentially with: (1) 2% sodium dodecyl sulfate in ultrapure water, (2) 80 mM sodium phosphate, 2 M NaCl, 0.2% Tween 20, pH 7.8, (3) 100 mM sodium carbonate, (4) 80% isopropyl alcohol in ultrapure water, (5) ultrapure water, and (6) 50 mM ammonium bicarbonate. Peptides were released by digestion with PNGase F (Promega, Madison, WI) overnight at 37 °C with vortexing. Deglycosylated peptides were cleaned and desalted using the SP2 procedure (73). Peptide Retention Time Calibration (PRTC) Mixture (ThermoFisher Scientific) was added to each sample at a final concentration of 2 nM to enable retention time calibration and assessment of instrument performance throughout the acquisition. A “pooled QC” mixture was generated by combining equal portions of each sample. Individual samples were queued in a randomized order within a technical injection series with two technical replicates each. Pooled QC samples were analyzed at the beginning and end of each technical injection series. Samples were analyzed by liquid chromatography tandem mass spectrometry (LC-MS/MS) using a Dionex UltiMate 3000 RSLCnano system in-line with an Orbitrap Fusion Lumos (ThermoFisher Scientific), and data were analysed with Proteome Discoverer 2.3 and Skyline (38). Normalized quantitation ratios were determined by comparisons of each sample type to the pooled QC.

### Immunofluorescence microscopy

T cells were adhered onto poly-L-lysine-coated glass coverslips as described in (74) and permeabilized in 0.5% saponin in PBS + 3% BSA + 880 uM Ca^2+^ + 490 uM Mg^2+^. Permeabilized cells were incubated with primary antibodies, washed, and then incubated with secondary antibodies conjugated to a fluorophore. 4’,6-diamidino-2-phenylindole (DAPI) was added to the final PBS wash at 1 ug/ml to stain DNA. Superresolution microscopy was performed on a Nikon Structured-Illumination Microscope (N-SIM; Nikon) and NIS-Elements AR imaging and 3D reconstruction software (v. 5.11) (Nikon Instruments Inc, Melville, NY). Images were taken using a Nikon 100X Oil-immersion lens (CFI Apo SR TIRF 1.49 NA) and an Andor iXon+897 EMCCD camera. Confocal microscopy was performed on a Nikon Eclipse Ti2 microscope equipped with a W1 Spinning Disc, Orca Flash CMOS camera, and 100X oil-immersion objective (CFI Plan Apo λ 1.49 NA), and NIS-Elements AR imaging and 3D reconstruction software (v. 6.0).

### Flow cytometry

Cells were incubated with primary antibodies in 1% bovine serum albumin (BSA) in DMEM -phenol red for 30 min on ice, washed, and incubated with secondary antibodies. Flow cytometry was performed using an LSRII flow cytometer (BD Biosciences, San Jose, CA). Data was analyzed using FlowJo analysis software (v. 10.7, BD Biosciences). Infected cells were selectively analyzed by GFP+ gating. Non-viable cells were excluded from all flow cytometric analyses.

### Immunoblotting

Cell lysates were prepared using 1% Triton Tx-100 lysis buffer supplemented with 62.5 U/ml Benzonase and 174 μg/ml phenylmethylsulfonyl fluoride (PMSF) followed by the addition of an equal volume of 2% sodium dodecyl sulfate (SDS) and 100 mM Tris-HCl (pH 7.4). Lysates were normalized to total protein concentration as determined by bicinchronic acid (BCA) assay (Pierce, Rockford, IL). Lysates were resolved by SDS-PAGE and transferred to BA-85 nitrocellulose membrane (Cytivia, Marlborough, MA). Membranes were incubated with primary antibodies, followed by HRP-conjugated secondary antibody (BioRad Laboratories, Hercules, CA). Bands were visualized using SuperSignal West Pico reagent (ThermoFisher Scientific) imaged with an Azure c600 gel documentation system and quantified using AzureSpot software (v2.2.167) (Azure Biosystems, Dublin, CA).

### Inhibition of protein degradation

HHV6A-infected or uninfected JJhan cells were incubated in RPMI in 50 mM NH4Cl, 100 nM MG-132, or a DMSO control for 19 hours. Cells were then divided for use in immunofluorescence microscopy or immunoblot analysis.

### qRT-PCR

Cells were lysed in TRIzol (Invitrogen, Carlsbad, CA) according to the manufacturer’s instructions. Approximately 2 ug of RNA was treated with an AccuRT Genomic DNA Removal Kit (Applied Biological Materials, Richmond, BC, Canada) prior to cDNA synthesis using the 5X All-In-One Master mix (Applied Biological Materials), both according to the manufacturer’s instructions. qPCR was performed using a BioRad CFX96 Real-Time System (BioRad Laboratories, Hercules, CA) and data analyzed using BioRad CFX Maestro (v.4.1.2433.1219). CD45 RNA levels were normalized to a cellular control for each sample (28S rRNA). Primer sets are listed in Table 1. All qPCR reactions were run in technical triplicate with corresponding no template and -RT controls, which did not exceed background levels. The delta-delta Ct method was used to calculate the relative abundance of CD45 cDNA from HHV6A-infected JJhan cells and 293T cells relative to the uninfected JJhan cell control.

**Table 1.**
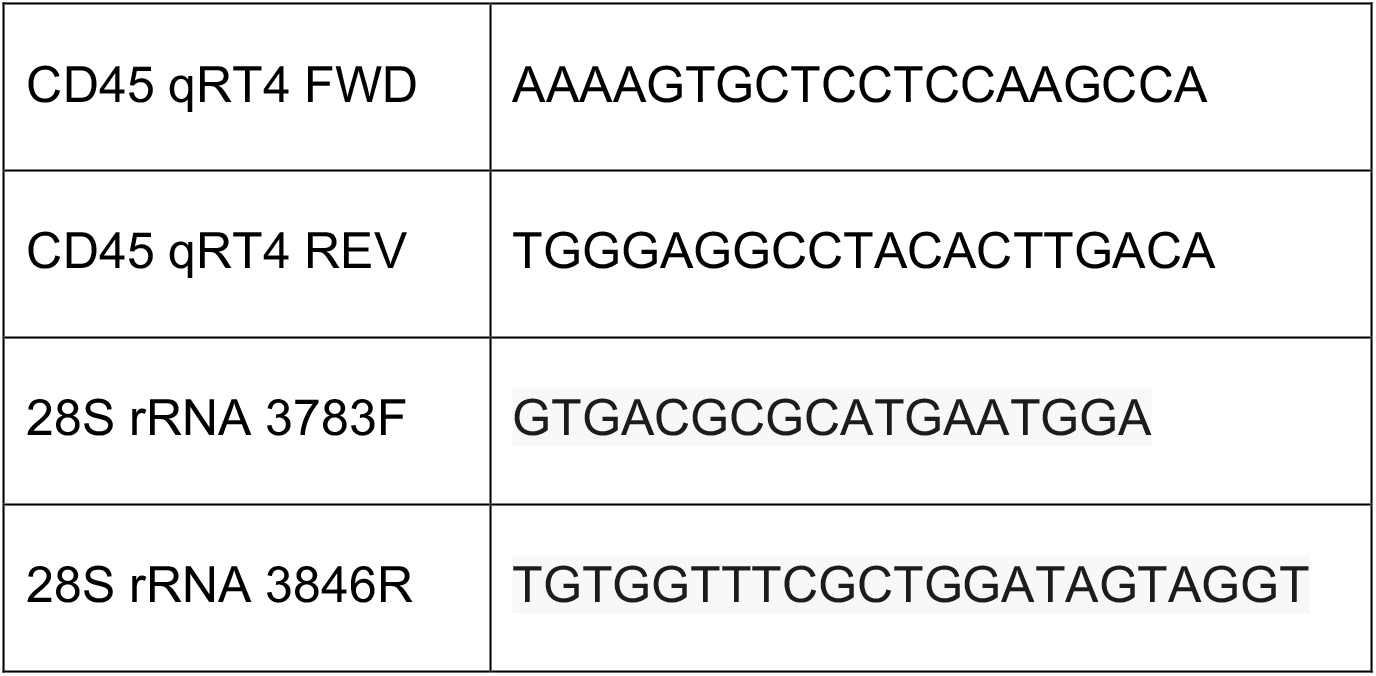
Primers used in this study.

### Statistical analysis

Statistical analysis was done using GraphPad 8 (v.8.4.3). One-way ANOVA analysis with Tukey’s multiple comparisons test was used to compare the abundance of CD45 mRNA 293T cells, uninfected JJhan cells, and HHV6A-infected T cells. **** < 0.0001

## Acknowledgements

The authors thank Rebekah Mokry for helpful discussion, Drs. Benedikt Kaufer, Arthur Weiss, Vera Tarakanova, Ken Brockman, and Scott Terhune for generous provision of reagents and helpful discussion. We acknowledge funding from National Institutes of Health under Award Numbers AI123745 to AWH and RLG, HL134010 to RLG, and the American Heart Association 20PRE35200049 to LBL. LBL is a member of the MCW Medical Scientist Training Program, partially supported by NIH T32 GM080202.

