## Supplemental figures for "The roseoloviruses downregulate the protein tyrosine phosphatase PTPRC (CD45)"

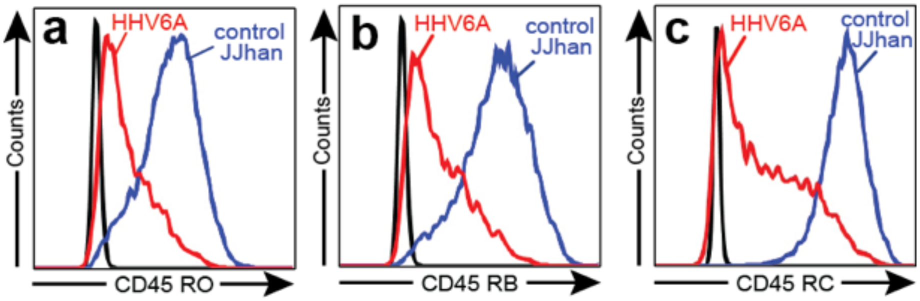


**Figure S1. All isoforms of CD45 are downregulated in HHV6A-infected cells.** Flow cytometry analysis of surface-expressed CD45 isoforms in HHV6A-infected JJhan cells (red) and uninfected JJhan cells (blue). Live cells were labeled with an antibody directed against CD45 isoforms (a) CD45RO, (b) CD45RB, or (c) CD45RC. Infected cells were gated on as GFP+. No primary antibody is shown in (black).

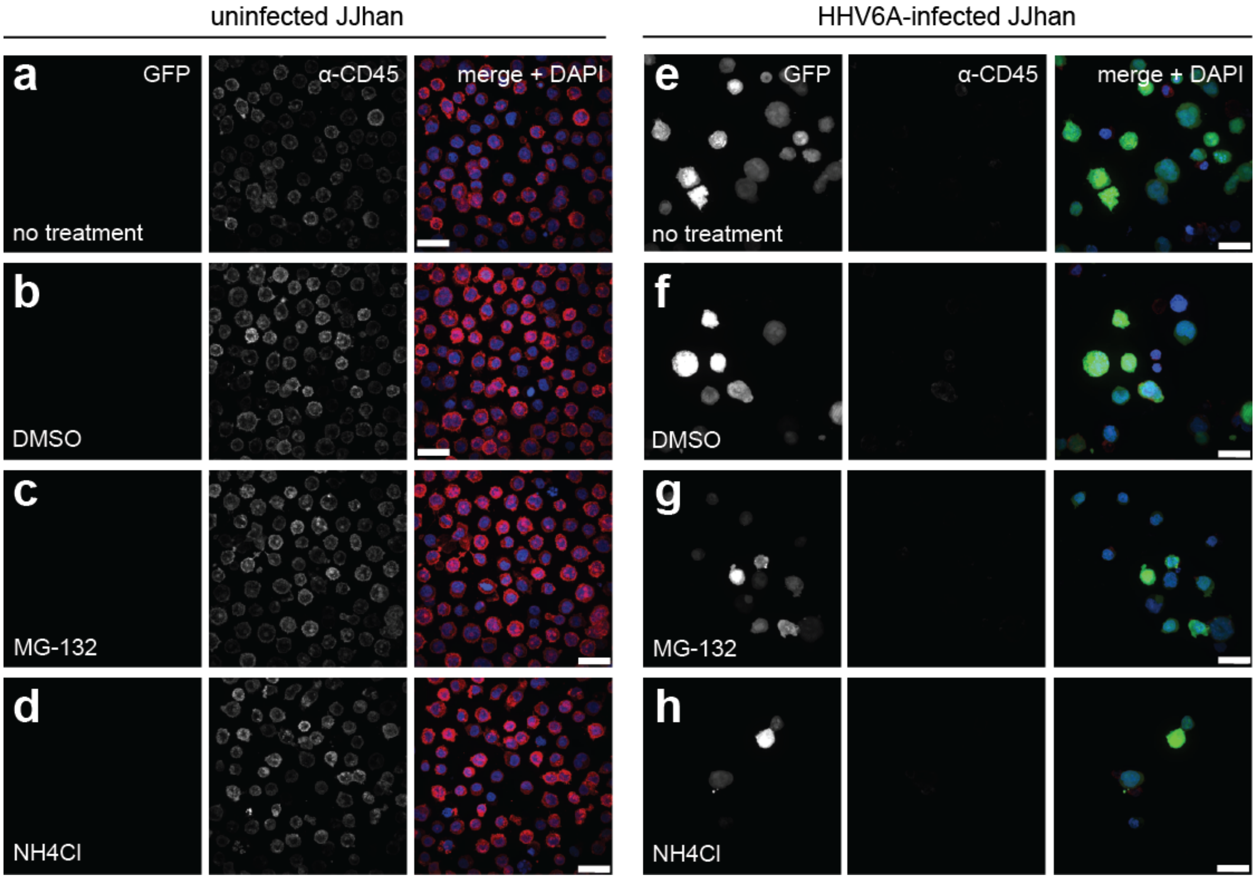


**Figure S2. Surface expression of CD45 is not restored in HHV6A-infected cells treated with either MG-132 or $NH_4Cl$.** Confocal immunofluorescence microscopy of (a-d) uninfected JJhan cells and (e-h) HHV6A-infected JJhan cells. HHV6A-infected cells express soluble GFP. Cells were either untreated (a, e), treated with a DMSO vehicle (b, f), treated with MG-132 (c, g), or treated with $NH_4Cl$ (d, h) for 19 hours before being fixed, permeabilized, and labeled with an antibody directed against CD45. Images were taken at 100X magnification and are shown as maximum intensity projections. Scale bar = 20 μm.
